# Hybracter: Enabling Scalable, Automated, Complete and Accurate Bacterial Genome Assemblies

**DOI:** 10.1101/2023.12.12.571215

**Authors:** George Bouras, Ghais Houtak, Ryan R. Wick, Vijini Mallawaarachchi, Michael J. Roach, Bhavya Papudeshi, Lousie M. Judd, Anna E. Sheppard, Robert A. Edwards, Sarah Vreugde

**Affiliations:** Adelaide Medical School, Faculty of Health and Medical Sciences, The University of Adelaide, Adelaide, Australia; The Department of Surgery - Otolaryngology Head and Neck Surgery, University of Adelaide and the Basil Hetzel Institute for Translational Health Research, Central Adelaide Local Health Network, South Australia, Australia; Department of Microbiology and Immunology, University of Melbourne at the Peter Doherty Institute for Infection and Immunity, Melbourne, Australia; Flinders Accelerator for Microbiome Exploration, College of Science and Engineering, Flinders University, Adelaide, Australia; Adelaide Centre for Epigenetics and South Australian Immunogenomics Cancer Institute, The University of Adelaide, Adelaide, Australia; School of Biological Sciences, The University of Adelaide, Adelaide, Australia

## Abstract

Improvements in the accuracy and availability of long-read sequencing mean that complete bacterial genomes are now routinely reconstructed using hybrid (i.e. short- and long-reads) assembly approaches. Complete genomes allow a deeper understanding of bacterial evolution and genomic variation beyond single nucleotide variants (SNVs). They are also crucial for identifying plasmids, which often carry medically significant antimicrobial resistance (AMR) genes. However, small plasmids are often missed or misassembled by long-read assembly algorithms. Here, we present Hybracter which allows for the fast, automatic, and scalable recovery of near-perfect complete bacterial genomes using a long-read first assembly approach. Hybracter can be run either as a hybrid assembler or as a long-read only assembler. We compared Hybracter to existing automated hybrid and long-read only assembly tools using a diverse panel of samples of varying levels of long-read accuracy with manually curated ground truth reference genomes. We demonstrate that Hybracter as a hybrid assembler is more accurate and faster than the existing gold standard automated hybrid assembler Unicycler. We also show that Hybracter with long-reads only is the most accurate long-read only assembler and is comparable to hybrid methods in accurately recovering small plasmids.

**Data Summary:**

1. Hybracter is developed using Python and Snakemake as a command-line software tool for Linux and MacOS systems.
2. Hybracter is freely available under an MIT License on GitHub (https://github.com/gbouras13/hybracter) and the documentation is available at Read the Docs (https://hybracter.readthedocs.io/en/latest/).
3. Hybracter is available to install via PyPI (https://pypi.org/project/hybracter/) and Bioconda (https://anaconda.org/bioconda/hybracter). A Docker/Singularity container is also available at https://quay.io/repository/gbouras13/hybracter.
4. All code used to benchmark Hybracter, including the reference genomes, is publicly available on GitHub (https://github.com/gbouras13/hybracter_benchmarking) with released DOI (https://zenodo.org/doi/10.5281/zenodo.10910108) available at Zenodo.
5. The subsampled FASTQ files used for benchmarking are publicly available at Zenodo with DOI (https://doi.org/10.5281/zenodo.10906937).
6. All super accuracy simplex ATCC FASTQ reads sequenced as a part of this study can be found under BioProject PRJNA1042815.
7. All *Hall* et al. fast accuracy simplex and super accuracy duplex ATCC FASTQ read files (prior to subsampling) can be found in the SRA under BioProject PRJNA1087001.
8. All raw *Lermaniaux* et al. FASTQ read files and genomes (prior to subsampling) can be found in the SRA under BioProject PRJNA1020811.
9. All *Staphylococcus aureus* JKD6159 FASTQ read files and genomes can be found under BioProject PRJNA50759.
10. All *Mycobacterium tuberculosis* H37R2 FASTQ read files and genomes can be found under BioProject PRJNA836783.
11. The complete list of BioSample accession numbers for each benchmarked sample can be found in Supplementary Table 1.
12. The benchmarking assembly output files are publicly available on Zenodo with DOI (https://doi.org/10.5281/zenodo.10906937).
13. All Pypolca benchmarking outputs and code are publicly available on Zenodo with DOI (https://zenodo.org/doi/10.5281/zenodo.10072192).

**Impact Statement:** Complete bacterial genome assembly using hybrid sequencing is a routine and vital part of bacterial genomics, especially for identification of mobile genetic elements and plasmids. As sequencing becomes cheaper, easier to access and more accurate, automated assembly methods are crucial. With Hybracter, we present a new long-read first automated assembly tool that is faster and more accurate than the widely-used Unicycler. Hybracter can be used both as a hybrid assembler and with long-reads only. Additionally, it solves the problems of long-read assemblers struggling with small plasmids, with plasmid recovery from long-reads only performing on par with hybrid methods. Hybracter can natively exploit the parallelisation of high-performance computing (HPC) clusters and cloud-based environments, enabling users to assemble hundreds or thousands of genomes with one line of code. Hybracter is available freely as source code on GitHub, via Bioconda or PyPi.

## Introduction

Reconstructing complete bacterial genomes using *de novo* assembly methods had been considered too costly and time-consuming to be widely recommended in most cases, even as recently as 2015 ^1^. This was due to the reliance on short-read sequencing technologies, which does not allow for reconstructing regions with repeats and extremely high GC content ^2^. However, since then, advances in long-read sequencing technologies have allowed for the automatic construction of complete genomes using hybrid assembly approaches. Originally, this involved starting with a short-read assembly followed by scaffolding the repetitive and difficult to resolve regions with long-reads ^3,4^. This approach was implemented in the command-line tool Unicycler, which remains the most popular tool for generating complete bacterial genome assemblies ^5^. As long-read sequencing has improved in accuracy and availability, with the latest Oxford Nanopore Technologies reads recently reaching Q20 (99%+) median accuracy, a long-read first assembly approach supplemented by short-read polishing has recently been favoured for recovering accurate complete genomes. Long-read-first approaches provide greater accuracy and contiguity than short-read-first approaches in difficult regions ^6–11^. The current gold standard manual assembly tool Trycycler even allows for the potential recovery of perfect genome assemblies ^7^. However, Trycycler requires significant microbial bioinformatics expertise and involves manual decision making, creating a significant barrier to useability, scalability and automation ^12^.

Several tools exist that generate automated long-read first genome assemblies, such as MicroPIPE ^13^, ASA3P ^14^, Bactopia ^15^ and Dragonflye ^16^. However, these tools do not consider factors such as genome reorientation ^17^ and recent polishing best-practices ^18^, and often contain the assembly workflow as a sub-module within a more expansive end-to-end pipeline. Additionally, none of the existing tools consider the targeted recovery of plasmids. As long-read assemblers struggle particularly with small plasmids, this leads to incorrectly recovered or missing plasmids in bacterial assemblies ^19^.

We introduce Hybracter, a new command-line tool for automated near-perfect long-read-first complete bacterial genome assembly. It implements a comprehensive and flexible workflow allowing for long-read assembly polished with long and short-reads (with subcommand ‘hybracter hybrid’ for one or more samples and subcommand ‘hybracter hybrid-single’ for a single sample) or long-read only assembly polished with long-reads (with subcommand ‘hybracter long’ for one or more samples and subcommand ‘hybracter long-single’ for a single sample) (Table 1). For ease of use and familiarity, Hybracter has been designed with a command-line interface containing parameters similar to Unicycler. Additionally, thanks to its Snakemake ^20^ and Snaketool ^21^ implementation, Hybracter seamlessly scales from a single isolate to hundreds or thousands of genomes with high computational efficiency and supports deployment on HPC clusters and cloud-based environments.

**Table 1.** Summary of the 4 Primary Hybracter Commands.

| Command | Input | Number of Samples | Description | Workflow Elements Included by Default (From Figure 1) |
| --- | --- | --- | --- | --- |
| **hybracter hybrid** | 5 column csv sample sheet specified with '--input' containing:<br>sample name<br>long-read FASTQ path,<br>estimated chromosome length<br>R1 short-read FASTQ path<br>R2 short-read FASTQ path | 1+ | Long-read first assembly followed by long then short-read polishing for multiple isolates. Snakemake implementation ensures efficient use of available resources | A, B, C, D, E, F, G, H |
| **hybracter hybrid-single** | sample name (-s)<br>long-read FASTQ path (-l)<br>estimated chromosome length (-c)<br>R1 short-read FASTQ path (-1)<br>R2 short-read FASTQ path (-2) | 1 | Long-read first assembly followed by long then short-read polishing for a single isolate. Similar command line interface to Unicycler | A, B, C, D, E, F, G, H |
| **hybracter long** | 3 column csv sample sheet specified with '--input' containing:<br>sample name<br>long-read FASTQ path,<br>estimated chromosome length | 1+ | Long-read first assembly followed by long-read polishing for multiple isolates. Snakemake implementation ensures efficient use of available resources | A (no fastp), B, C, D, E, G, H |
| **hybracter long-single** | sample name (-s)<br>long-read FASTQ path (-l)<br>estimated chromosome length (-c) | 1 | Long-read first assembly followed by long-read polishing on a single isolate. | A (no fastp), B, C, D, E, G, H |

## Methods

### Assembly Workflow

Hybracter implements a long-read-first automated assembly workflow based on current best practices ^12^. The main subcommands available in Hybracter can be found in Table 1 and the workflow is outlined in Figure 1. Hybracter begins with long-reads for all subcommands, and uses short-reads for polishing for ‘Hybracter hybrid’ and ‘Hybracter hybrid-single’ subcommands.

**Figure 1:**
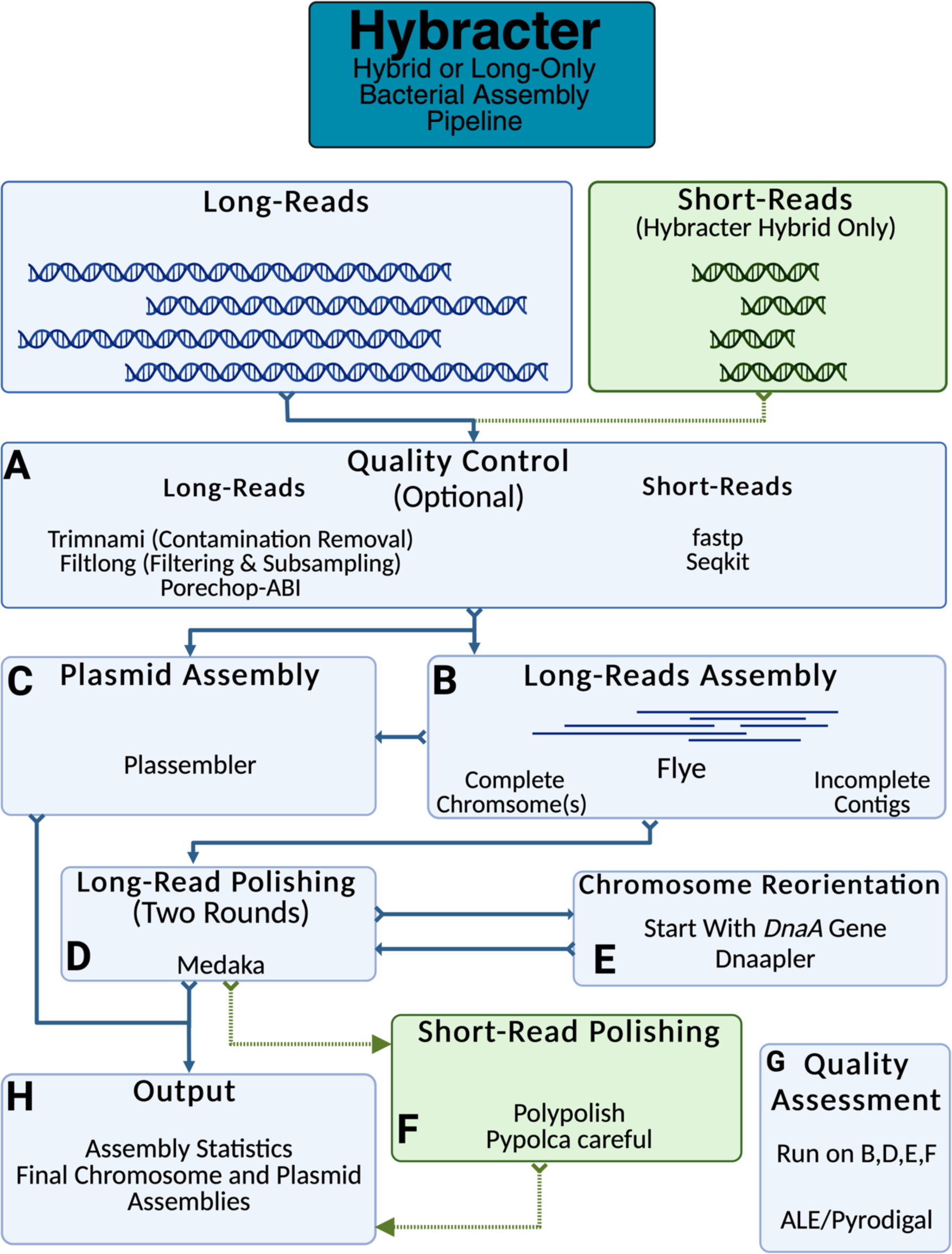
Outline of the Hybracter workflow.

First, long-read input FASTQs are input long read sets are filtered and subsampled to a depth of 100x with Filtlong^22^, which prioritises the longest and highest quality reads, outperforming random subsampling (See Supplementary Table 11). The reads also have adapters trimmed using Porechop_ABI^23^, with optional contaminant removal against a host genome using modules from Trimnami (e.g. if the bacteria has been isolated from a host) ^24^. Quality control of short-read input FASTQs is performed with fastp ^25^ (Fig 1A). The estimated depth of the short-reads is determined using Seqkit^26^.

Long-reads are then assembled with Flye ^27^. If at least 1 contig is recovered above the cut-off ‘-c’ chromosome length specified by the user for the sample, that sample will be denoted as ‘complete’. All such contigs will then be marked as chromosomes and kept for downstream polishing and reorientation if marked as circular by Flye. If zero contigs are above the cut-off chromosome length, the assembly will be denoted as ‘incomplete’, and all contigs will be kept for downstream polishing (Fig 1B).

For all complete samples, targeted plasmid assembly is then conducted using Plassembler ^28^ (Fig 1C). All samples (i.e. complete and incomplete) are then polished once with Medaka^29^, which can be turned off using ‘--no_medaka’ (Fig 1D). It is recommended to turn off Medaka using ‘--no_medaka’ for highly accurate Q20+ read sets where Medaka has been shown to introduce false positive changes^11^. For all complete samples only, chromosome(s) marked as circular by Flye will then be reoriented to begin with the dnaA chromosomal replication initiator gene using Dnaapler^30^. These reoriented chromosomes are then polished for a second time with Medaka to ensure the sequence around the original chromosome breakpoint is polished.

If the user has provided short-reads with Hybracter hybrid, all samples’ assemblies (complete and incomplete) are then polished with Polypolish ^18^ followed by Pypolca ^31,32^ (Fig 1F). The exact parameters depend on the depth of short-read sequencing^31^. If the estimated short-read coverage is below 5x, only Polypolish with ‘--careful’ is run, as Pypolca can rarely introduce false positive errors at low depths. If the estimated short-read coverage is between 5-25x, Polypolish with --careful parameter is run followed by Pypolca with --careful parameter. Above 25x coverage, Polypolish with default parameters followed by Pypolca with --careful is run. This is because Pypolca --careful has been shown to be the best polisher at depths above 5x, and because Polypolish is able to fix potential errors in repeats Pypolca may miss. By default, the last short-read polishing round is chosen as the final assembly. Alternatively, users can choose the highest scoring polishing round according to the reference-free ALE ^33^ score.

If only long-reads are available (Hybracter long), the mean coding sequence (CDS) length is calculated for each assembly using Pyrodigal ^34,35^, with larger mean CDS lengths indicating a better quality assembly. The polishing round with the highest mean CDS length is chosen as the final assembly (Fig 1G).

For each sample, a final output assembly FASTA file is created, along with per contig and overall summary statistic TSV files, as well as separate chromosome and plasmid FASTA files for samples denoted as complete (Fig 1H). An overall ‘hybracter_summary.tsv’ file is also generated, which summarises outputs for all samples. All main output files are explained in more detail in Table 2. All the main outputs can be found in the ‘FINAL_OUTPUT’ subdirectory, while all other intermediate output files are available in other subdirectories for users who would like extra information about their assemblies, including all assembly assessments, comparisons of all changes introduced by polishing, and Flye and Plassembler output summaries. A full list of these supplementary outputs can be found in Hybracter’s Documentation (https://hybracter.readthedocs.io/en/latest/output/).

**Table 2.**
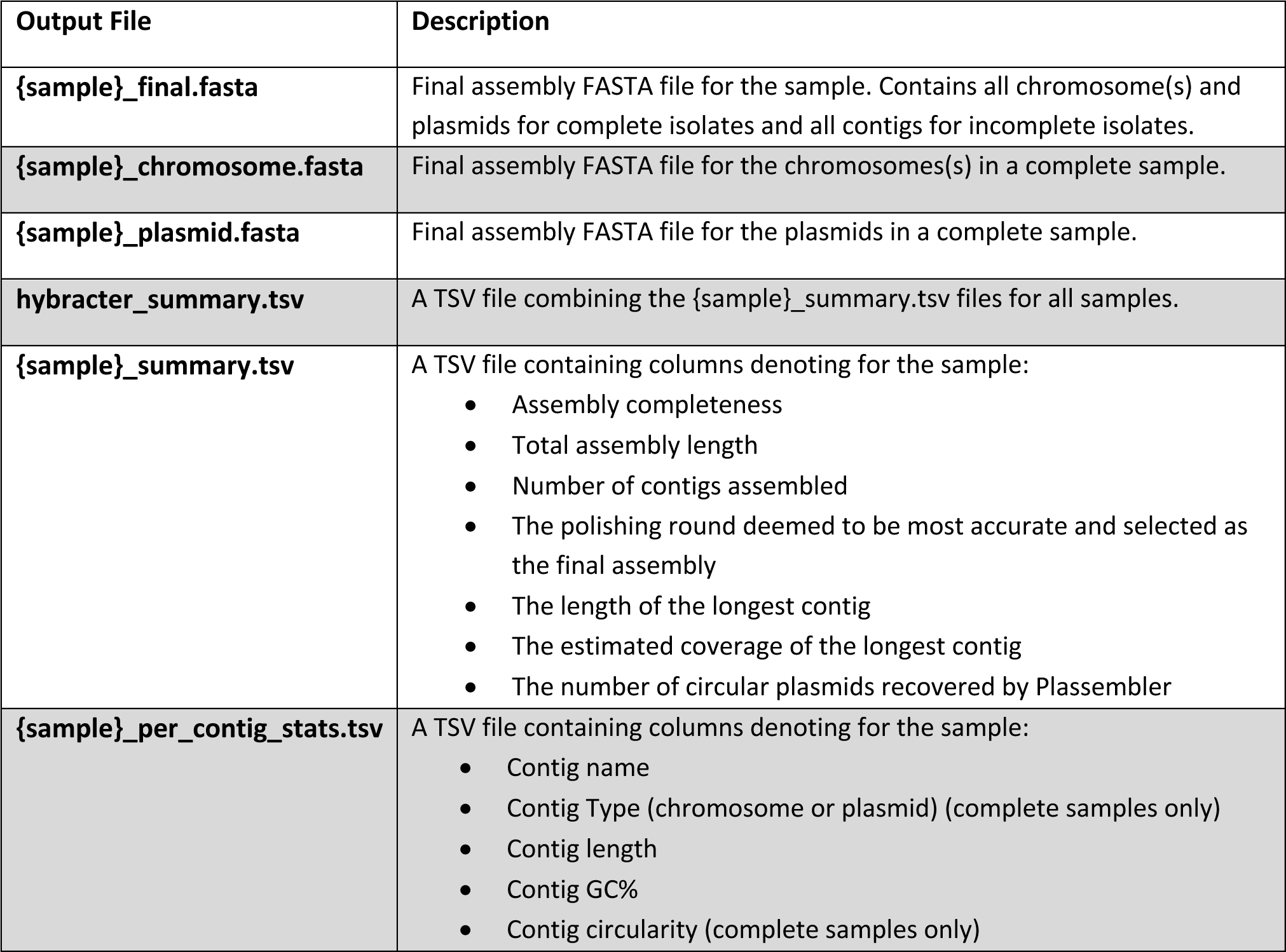
Description of the Primary Hybracter Output Files.

### Tool Selection

Tools were selected for inclusion in Hybracter either based on benchmarking from the literature, or they were specifically developed for inclusion in Hybracter. Flye ^27^ was chosen as the long-read assembler because it is more accurate for bacterial genome assembly than other long-read assemblers with comparable runtimes, such as Raven ^36^, Redbean ^37^ and Miniasm ^38^, while being dramatically faster than the comparably accurate Canu ^6,39^. Medaka ^29^ was chosen as the long-read polisher because of its ability to improve assembly continuity in addition to accuracy ^12,40^. The benchmarking results of this study also emphasise that it is particularly good at fixing insertion and deletion (InDel) errors, which cause problematic frameshifts and frequently lead to fractured or truncated gene predictions. However, it should be re-iterated that for modern Q20+ datasets, Medaka may introduce errors^11^ and should not be used (using --no_medaka with Hybracter). Polypolish and Pypolca in various combinations depending on short-read depth were selected as short-read polishers, as these have been shown to achieve the highest performance with the lowest chance of introducing errors when used in combination ^31^.

We developed three standalone programs included in Hybracter. These are Dnaapler^30^, Plassembler^28^ and Pypolca^31^. Dnaapler was developed to ensure the chromosome(s) identified by Hybracter are reoriented to consistently begin with the dnaA chromosomal replication initiator gene. Full implementation details can be found in the manuscript, with expanded functionality beyond this use case^30^. Plassembler was developed to improve the runtime and accuracy when assembling plasmids in bacterial isolates. Full implementation details can be found in the manuscript for hybrid mode ^28^. Hybracter long utilises Plassembler containing a post-publication improvement for long-reads only (‘Plassembler long’) released in v1.3. Plassembler long assembles plasmids from only long-reads by treating long-reads as both short-reads and long-reads. Plassembler long does this by utilising Unicycler in its pipeline to create a de Bruijn graph-based assembly, treating the long-reads as unpaired single-end reads, which are then scaffolded with the same long-read set.

The third tool is Pypolca^31,32^. Pypolca is a Python re-implementation of the POLCA short-read genome polisher, originally created specifically for inclusion in Hybracter and with an almost identical output format and performance. Compared to POLCA, Pypolca features improved useability with a simplified command line interface, allows the user to specify an output directory and introduces a ‘--careful’ parameter. The performance of Pypolca, and particularly Pypolca with the --careful parameter, are described in the manuscript^31^.

### Benchmarking

To compare Hybracter’s functionality and performance, we benchmarked its performance against other software tools. We focused on the most popular state-of-the-art assembly tools for automated hybrid and long only bacterial genome assemblies. All code to replicate these analyses can be found at the repository (https://github.com/gbouras13/hybracter_benchmarking). All programs and dependency versions used for benchmarking can be found in Supplementary Table 4. For the hybrid tools, we chose Unicycler and Dragonflye with both long-read and short-read polishing (denoted ‘Dragonflye hybrid’). Dragonflye was chosen as it is a popular long-read first assembly pipeline ^16^. Both tools were run using default parameters. By default, Dragonflye conducts a long-read assembly with Flye that is polished by Racon^41^ followed by Polypolish. For the long-read only tool, we chose Dragonflye with long-read Racon based polishing only (denoted ‘Dragonflye long’).

We used 30 samples for benchmarking, representing genomes from a variety of Gram-negative and Gram-positive bacteria. We chose these samples as they have real hybrid read sets in combination with manually curated genome assemblies produced using either Trycycler or Bact-builder^42^—a consensus-building pipeline based on Trycycler. These samples came from 5 different studies below. We used the published genomes from these studies as representatives of the ‘ground truth’ for these samples. Where read coverage exceeded 100x samples were subsampled to approximately 100x coverage of the approximate genome size with Rasusa v0.7.0^43^, as this better reflects more realistic read depth of real life isolate sequencing. Nanoq v0.10.0^44^ was used to generate quality control statistics for the subsampled long-read sets. Four isolates did not have 100x long-read coverage — the entire long-read set was used instead. A full summary table of the read lengths, quality, Nanopore kit and base-calling models used in these studies can be found in Supplementary Table 2.

Hybracter v0.7.0 was used to conduct benchmarking. Medaka long-read polishing was used for all samples except the 5 ATCC super-accuracy model basecalled duplex read samples, where ‘--no_medaka’ was used.

These samples contained varying levels of long-read quality (reflecting improvements in Oxford Nanopore Technologies long-read technology), with the median Q score of long-read sets ranging from 10.6 to 26.8. The five studies are:

1. Five ATCC strain isolates (ATCC-10708 *Salmonella enterica*, ATCC-17802 *Vibrio paragaemolyticus*, ATCC-25922 *Escherichia coli*, ATCC-33560 *Campylobacter jejuni* and ATCC-BAA-679 *Listeria monocytogenes*) with R10 chemistry super-accuracy model basecalled simplex long-reads made available as a part of this study.
2. The same 5 ATCC isolates with R10 chemistry fast model basecalled long-reads, and R10 chemistry super-accuracy model basecalled duplex long-reads from *Hall* et al.^45^
3. Twelve diverse carbapenemase-producing Gram-negative bacteria from *Lerminiaux* et al.^9^
4. *Staphylococcus aureus* JKD6159 sequenced with both R9 and R10 chemistry long-read sets from *Wick* et al.^46^
5. *Mycobacterium tuberculosis* HR37v from *Chitale* et al.^42^

The full details for each individual isolate used can be found in Supplementary Tables 1 and 2.

### Chromosome Accuracy

The assembly accuracy of the chromosomes recovered by each benchmarked tool was compared using Dnadiff v1.3 packaged with MUMmer v3.23^47^. Comparisons were performed on the largest assembled contig (denoted as the chromosome) by each method, other than for ATCC-17802 *Vibrio parahaemolyticus*, where the two largest contigs were chosen as it has two chromosomes.

### Plasmid Recovery Performance and Accuracy

Plasmid recovery performance for each tool was compared using the following methodology. Summary statistics are presented in Table 4. See Supplementary Table 7 for a full sample-by-sample analysis. All samples were analysed using the 4-step approach outlined below using summary length and GC% statistics for all contigs and the output of Dnadiff v1.3 comparisons generated for each sample and tool combination against the reference genome plasmids:

1. The number of circularised plasmid contigs recovered for each isolate was compared to the reference genome. If the tool recovered a circularised contig homologous to that in the reference, it was denoted as completely recovered. Specifically, a contig was denoted as completely recovered if it had a genome length within 250bp of the reference plasmid, a GC% within 0.1% of the reference plasmid and whether the Total Query Bases covered was within 250bp of the Total Reference Bases from Dnadiff. For Dragonflye assemblies, some plasmids were duplicated or multiplicated due to known issues with the long-read first assembly approach for small plasmids ^6,19,48^. Any circularised contigs that were multiplicated compared to the reference plasmid were therefore denoted as misassembled.
2. For additional circularised contigs not found in the reference recovered, these were tested for homology with NCBI nt database using the web version of blastn^48^. If there was a hit to a plasmid, the Plassembler output within Hybracter was checked for whether the contig had a Mash hit (i.e. a Mash distance of 0.2 or lower) to plasmids in the PLSDB^49^. If there was a hit, the contig was denoted as an additional recovered plasmid. There were 2 in total (see Supplementary Table 7 and supplementary data).
3. Plasmids with contigs that were either not circularised but homologous to a reference plasmid, or circularised but incomplete (failing the genome length and Dnadiff criteria in 1.) were denoted as partially recovered or misassembled.
4. Reference plasmids without any homologous contigs in the assembly were denoted as missed.

Additional non-circular contigs that had no homology with reference plasmids and were not identified as plasmids in step 2 were analysed on a contig-by-contig basis and denoted as additional non-plasmid contigs (see Supplementary Table 7 for contig-by-contig analysis details).

### Runtime Performance Comparison

To compare the performance of Hybracter, we compared wall-clock runtime consumption on a machine with an Intel^®^ Core™ i9-13900 CPU @ 5.60 GHz on a machine running Ubuntu 20.04.6 LTS with a total of 32 available threads (24 total cores). We ran all tools with 8 and 16 threads and with 32 GB of memory to provide runtime metrics comparable to commonly available consumer hardware. Hybracter hybrid and long were run with ‘hybracter hybrid-single’ and ‘hybracter long-single’ for each isolate to generate a comparable per sample runtime for comparison with the other tools. The summary results are available in Table 5 and the detailed results for each specific tool and thread combination are found in Supplementary Table 8.

### Depth Analysis

To assess the effect of long-read depth on assembly accuracy, we chose *Lerminiaux* Isolate B (*Enterobacter cloacae*) and subsampled the long-read depth at each interval of 5x from 10x to 100x estimated genome size. All 5 tools were run on these read sets. Where a complete chromosome was assembled, Dnadiff (as described above) was used to compare the chromosome assembly to the reference.

### Sequencing

DNA extraction was performed with the DNeasy Blood and Tissue kit (Qiagen). Illumina library preparation was performed using Illumina DNA prep (Illumina Inc.) according to the manufacturer’s instructions. Short-read whole genome sequencing was performed an Illumina MiSeq with a 250bp PE kit. Oxford Nanopore Technologies library preparation ligation sequencing library was prepared using the ONT SQK-NBD114-96 kit and the resultant library was sequenced using an R10.4.1 MinION flow cell (FLO-MIN114) on a MinION Mk1b device. Data was base-called with Super-Accuracy Basecalling (SUP) using the basecaller model dna_r10.4.1_e8.2_sup@v3.5.1.

### Pypolca Benchmarking

Pypolca v0.2.0 was benchmarked against POLCA (in MaSuRCA v4.1.0)^32^ using 18 isolates described above. These were all 12 *Lerminiaux* et al. isolates, the R10 JKD6159 isolate ^46^ and the 5 ATCC samples we sequenced as a part of this study. Benchmarking was conducted on an Intel® Core™ i7-10700K CPU @ 3.80 GHz on a machine running Ubuntu 20.04.6 LTS. All short read FASTQs used for benchmarking are identical to those used to benchmark Hybracter. The assemblies used for polishing were intermediate chromosome assemblies from Flye v2.9.2^50^ generated within Hybracter. The outputs from Pypolca and POLCA were compared using Dnadiff v1.3 packaged with MUMmer v3.23^47^ Overall, Pypolca and POLCA yielded extremely similar results. 16/18 assemblies were identical. ATCC 33560 had 2 Single Nucleotide Polymorphisms (SNPs) between Pypolca and POLCA and *Lerminiaux* Isolate I also had 2 SNPs.

## Results

### Chromosome Accuracy Performance

All tools recovered complete circular contigs for each chromosome. SNVs, small InDels (under 60 bps), and large InDels (over 60 bps) were compared as a measure of assembly accuracy. To account for differences in genomic size between isolates, SNVs and small InDel counts were normalised by genome length.

The summary results are presented in Table 3 and visualised in Figure 2. The detailed results for each tool and sample are presented in Supplementary Table 5. Of the hybrid tools, Dragonflye hybrid and Hybracter hybrid produced the fewest SNVs (both with median 0) followed by Unicycler (median 34). Hybracter hybrid produced the fewest InDels (median 0), followed by Dragonflye hybrid (median 2.5) and Unicycler (median 11). Hybracter hybrid also produced the fewest InDels plus SNVs (median 1), followed by Dragonflye hybrid (median 4.5) and Unicycler (median 57.5).

**Figure 2:**
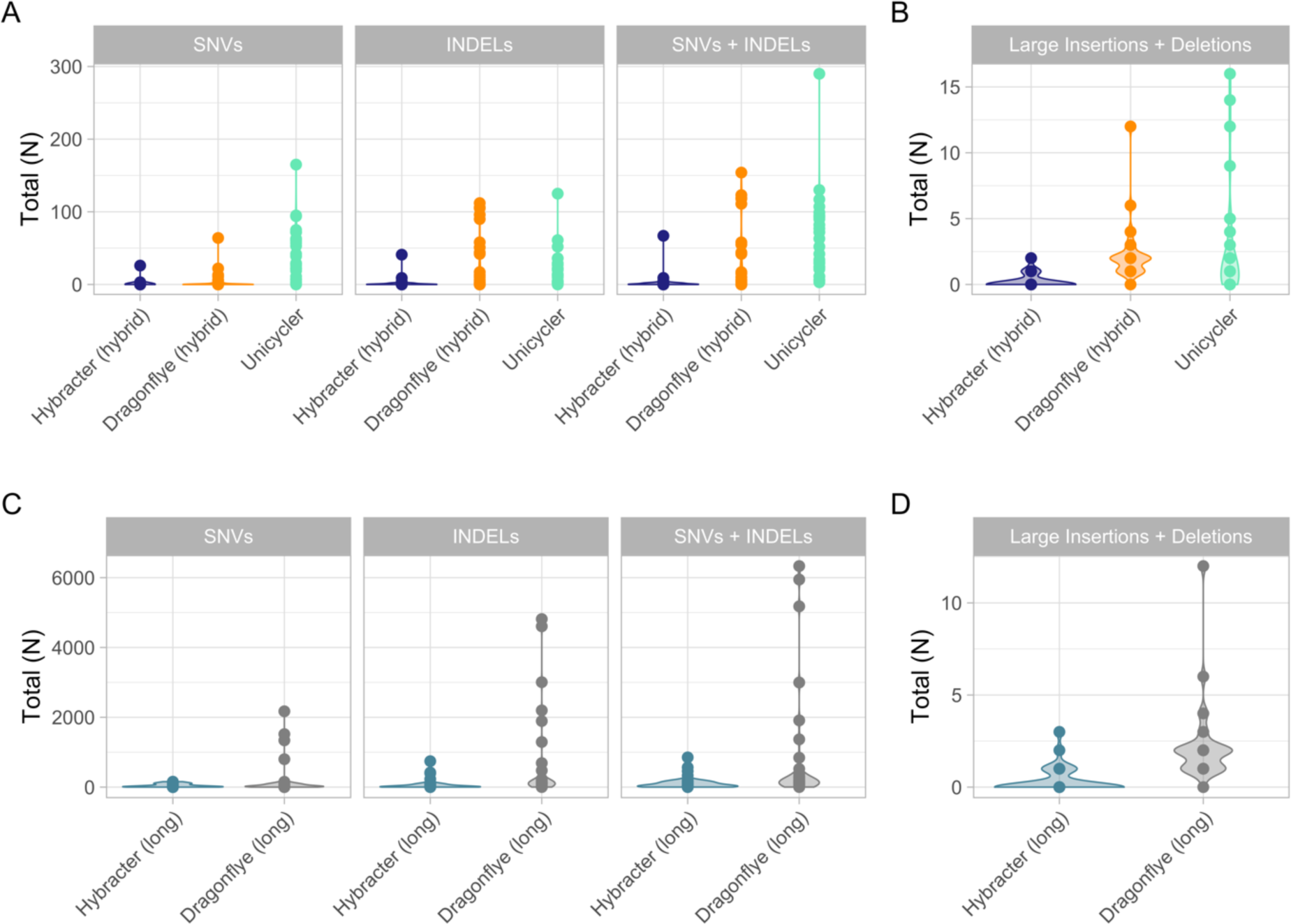
Comparison of the counts of single nucleotide variants (SNVs) and small (<60bp) insertions and deletions (InDels) (A) and the total number of large (>60bp) InDels (B) for the hybrid tools benchmarked (Hybracter hybrid in dark blue, Dragonflye hybrid in orange and Unicycler in green). The counts of SNVs and small InDels (C) and the total number of large InDels (D) for the long tools benchmarked (Hybracter long in light blue, Dragonflye long in grey) are also shown. All data presented is from the benchmarking output run with 8 threads.

**Table 3.** Small (<60bp) InDels, SNVs and large (>60bp) InDels of Chromosomes Assemblies for all Benchmarked Isolates.

| Tool | Type | Small InDels | SNVs | Small InDels + SNVs | Large InDels |
| --- | --- | --- | --- | --- | --- |
| **Hybracter hybrid** | Hybrid | Median = 0<br>Minimum = 0<br>Maximum = 41 | Median = 0<br>Minimum = 0<br>Maximum = 26 | Median = 1<br>Minimum = 0<br>Maximum = 67 | Total = 9<br>Median = 0<br>Minimum = 0<br>Maximum = 2 |
| **Dragonflye hybrid** | Hybrid | Median = 2.5<br>Minimum = 0<br>Maximum = 112 | Median = 0<br>Minimum = 0<br>Maximum = 64 | Median = 4.5<br>Minimum = 0<br>Maximum = 154 | Total = 70<br>Median = 2<br>Minimum = 0<br>Maximum = 12 |
| **Unicycler** | Hybrid | Median = 11<br>Minimum = 0<br>Maximum = 125 | Median = 34<br>Minimum = 0<br>Maximum = 165 | Median = 57.5<br>Minimum = 3<br>Maximum = 290 | Total = 87<br>Median = 1<br>Minimum = 0<br>Maximum = 16 |
| **Hybracter long** | Long | Median = 16<br>Minimum = 1<br>Maximum = 743 | Median = 21.5<br>Minimum = 0<br>Maximum = 156 | Median = 54<br>Minimum = 1<br>Maximum = 852 | Total = 11<br>Median = 1<br>Minimum = 0<br>Maximum = 3 |
| **Dragonflye long** | Long | Median = 125<br>Minimum = 2<br>Maximum = 4814 | Median = 34.5<br>Minimum = 0<br>Maximum = 2172 | Median = 170.5<br>Minimum = 2<br>Maximum = 6332 | Total = 68<br>Median = 2<br>Minimum = 0<br>Maximum = 12 |

Additionally, Hybracter hybrid showed superior performance in terms of large InDels, with a median of 0 and a total of 9 large InDels across the 30 samples, compared to 2 and 70 for Dragonflye hybrid, and 1 and 87 for Unicycler.

Overall, Hybracter hybrid produced the most accurate chromosome assemblies. For twelve isolates, Hybracter assembled a perfect chromosome (*Lerminiaux* et al.^9^ Isolates A, B, C, D, G, H, I, J, L, *S. aureus* JKD6159 with R10 chemistry and *L. monocytogenes* ATCC BAA-679 with simplex and duplex super-accuracy model basecalled reads).

Hybracter hybrid also produced several near-perfect assemblies (defined as <10 total SNVs plus InDels with no large insertions or deletions), including on some lower quality fast model basecalled reads (Supplementary Table 5).

Similar results were found in the long-read only tool comparison. Hybracter long produced the fewest SNVs (median 21.5) compared to Dragonflye long (median 34.5). Hybracter long consistently had far fewer small InDels (median 16) and large InDels (total 11 across 30 samples) compared to Dragonflye long (median 125 and total 68 respectively). No perfect chromosomes were assembled by either long-only tool, though Hybracter long did assemble three near-perfect chromosomes (*L. monocytogenes* ATCC BAA-679 with simplex and duplex super-accuracy model basecalled reads and ATCC 10708 *S. enterica* with duplex super-accuracy model basecalled reads) and several chromosomes with fewer than 50 total small InDels plus SNVs and 0 large InDels (*Lerminiaux* isolates A, G, H, L, J, and *S. aureus* JKD6159 with R10 chemistry, ATCC 10708 *S. enterica* with simplex super-accuracy model basecalled reads).

Overall, Hybracter long showed consistently worse performance than the hybrid tools Hybracter hybrid and Dragonflye hybrid tools (though not Unicycler) as measured by SNVs and small InDels. Combined with the lack of perfect assemblies even for duplex super-accuracy model basecalled read assemblies, this suggests the continuing utility of short-read polishing for the isolates surveyed.

### Plasmid Recovery Performance and Accuracy

Hybracter in both hybrid and long modes was superior at recovering plasmids compared to the other tools in the same class (Table 4). Hybracter hybrid was able to completely recover 65/69 possible plasmids (the other two were partially recovered), compared to 60/69 for Unicycler and only 44/69 for Dragonflye hybrid. Hybracter hybrid did not miss a single plasmid, while Unicycler missed 3/69 (all in Isolate E *Klebsiella pneumoniae* from *Lerminiaux* et al.) and Dragonflye hybrid completely missed 9/69. In terms of plasmid accuracy, Hybracter hybrid and Unicycler were similar in terms of SNVs plus small InDels, with medians of 1.62 and 2.02 per 100kbp respectively (Supplementary Table 9), while Hybracter hybrid produced fewer large InDels than Unicycler (44 vs 63 in total).

**Table 4.** The Total Number of Plasmids Recovered by Each Tool. There were 59 total reference plasmids in the 30 samples.

| Tool | Complete Plasmids Recovered | Total Plasmids Partially Recovered or Misassembled | Total Plasmids Missed | Additional Plasmids Recovered not in Reference | Samples with Additional Non-Plasmid Contigs Recovered |
| --- | --- | --- | --- | --- | --- |
| Hybracter hybrid | 65 | 4 | 0 | 2 | 10 |
| Unicycler | 60 | 6 | 3 | 1 | 2 |
| **Dragonflye**<br>**hybrid** | 44 | 16 | 9 | 1 | 10 |
| **Hybracter**<br>**long** | 60 | 5 | 4 | 2 | 3 |
| **Dragonflye**<br>**long** | 44 | 16 | 9 | 1 | 10 |

Interestingly, Hybracter long showed strong performance at recovering plasmids despite using only long-reads, completely recovering 60/69 plasmids and completely missing only 4/69. This performance was far superior to Dragonflye long (44/69 completely recovered, 9/69 missed). In terms of accuracy, both long tools were similar and unsurprisingly less accurate than the hybrid tools in terms of SNVs plus small InDels (medians of 8.74 per 100kbp for Hybracter long and 7.66 per 100kbp for Dragonflye long).

All five tools detected an additional 5411bp plasmid in *Lerminiaux* Isolate G not found in the reference sequence and Hybracter in both hybrid and long modes detected a further 2519bp small plasmid from this genome.

Hybracter hybrid recovers more plasmids than either Unicycler or Dragonflye because it uses a dedicated plasmid assembler, Plassembler. In addition, Hybracter long, using only long-reads had an identical complete plasmid recovery rate to Unicycler, which uses both long- and short-reads (60/69 for both). These results suggest that Hybracter long, by applying algorithms designed for short-reads on long-reads, largely solves the existing difficulties of recovering small plasmids from long-reads, at least on the benchmarking dataset of predominantly R10 Nanopore reads^19,51^. Even on the lower quality fast basecalled ATCC reads, Hybracter long performed well, with only one sample failing to produce a plasmid assembly similar to higher quality datasets (ATCC 10708 *S. enterica* – See Supplementary Tables 6 and 7).

Another notable result from Hybracter hybrid is that in 10/30 samples, it assembled additional non-plasmid contigs, which occurred in only 2/30 isolates for Unicycler. This is a limitation of Hybracter hybrid, as the extra sensitivity to recover plasmids comes with the cost of more false positive non-plasmid contigs that may be low-depth artefacts of sequencing. Hybracter has a ‘depth_filter’ parameter (defaulting to 0.25x of the chromosome depth) that filters out all non-circular putative plasmid contigs below this value.

It should be noted, however, that these contigs are not always an assembly artefact and can provide additional information regarding the quality control and similarity of short and long-read sets. In Plassembler implemented within Hybracter hybrid, the existence of such contigs is often indicative of mismatches between long- and short-read sets^28^, suggesting that there may be some heterogeneity between long- and short-reads in those samples.

### Runtime Performance Comparison

As shown in Table 5 and Figure 3, median wall-clock times with 8 threads for Dragonflye hybrid (4m34s) were smaller than Hybracter hybrid (15m03s), which were in turn smaller than Unicycler (50m25s). For the long-only tools, Dragonflye long (4m10s) was faster than Hybracter long (11m46s). Hybracter long was consistently slightly faster than Hybracter hybrid (Table 5).

**Figure 3:**
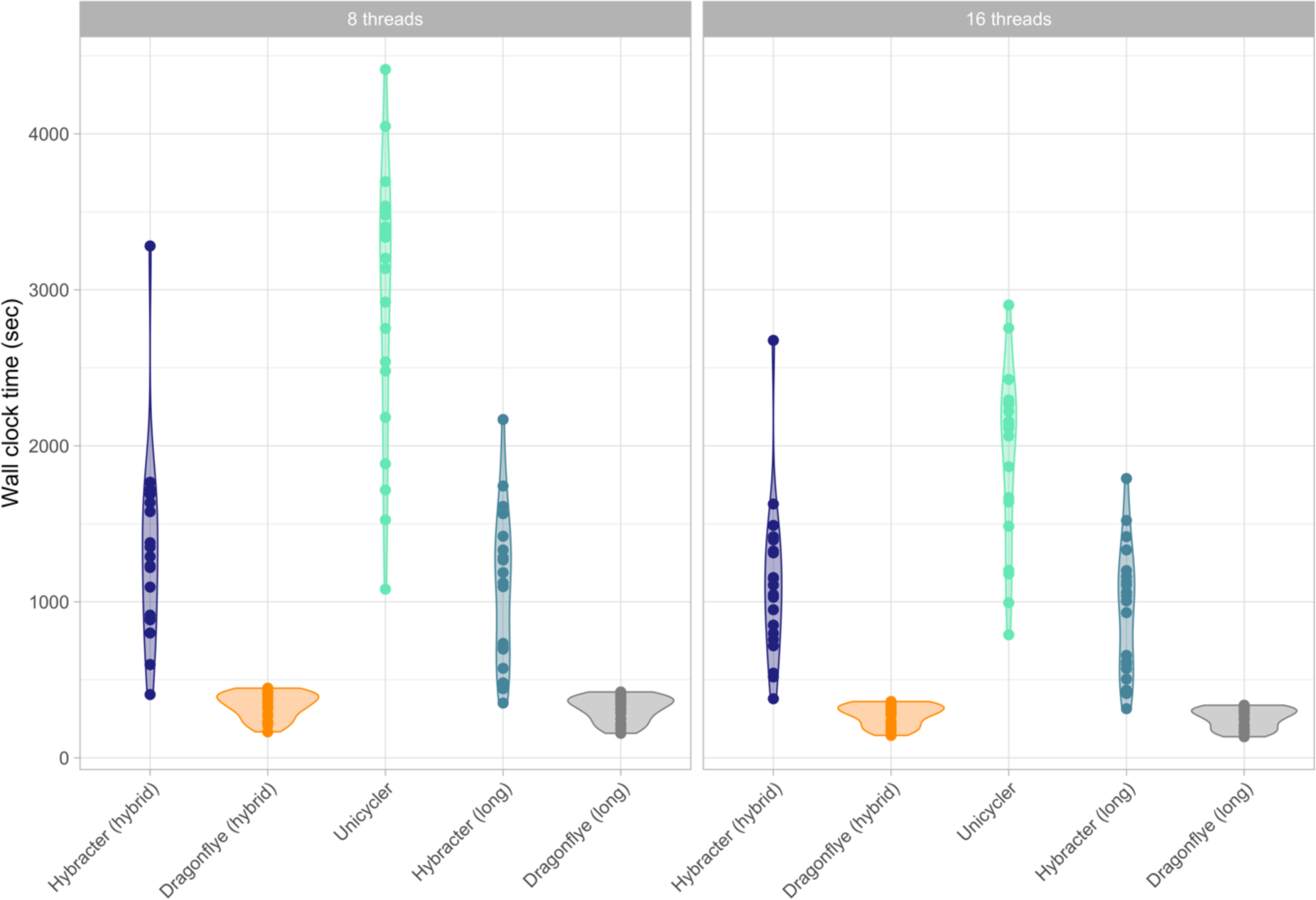
Comparison of wall-clock runtime (in seconds) of Hybracter hybrid, Dragonflye hybrid, Unicycler, Hybracter long and Dragonflye long when run with 8 and 16 threads.

**Table 5.**
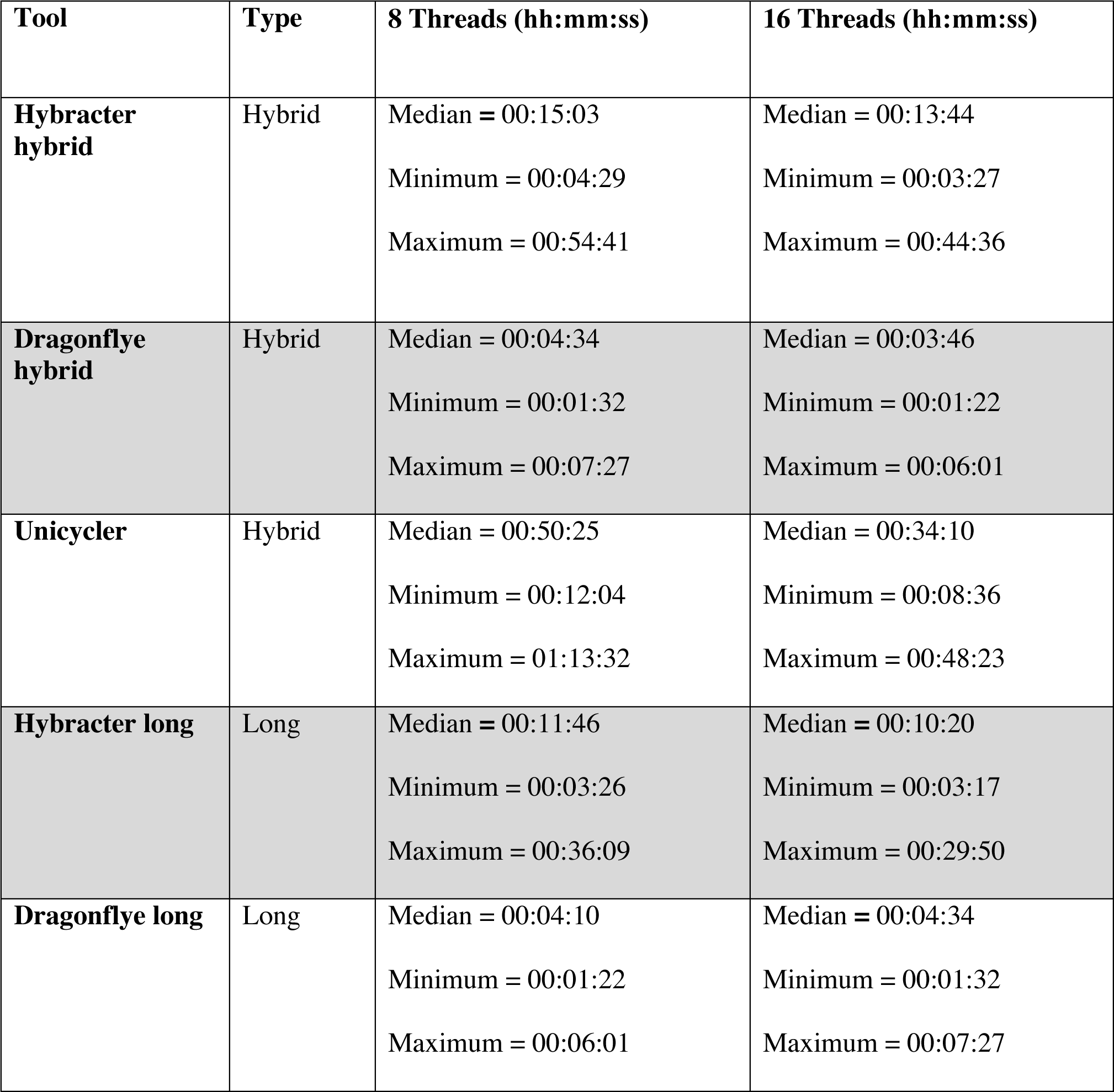
Wall-clock Runtime Summary Statistics for Each Tool.

| Tool | Type | 8 Threads (hh:mm:ss) | 16 Threads (hh:mm:ss) |
| --- | --- | --- | --- |
| **Hybracter hybrid** | Hybrid | Median = 00:15:03<br>Minimum = 00:04:29<br>Maximum = 00:54:41 | Median = 00:13:44<br>Minimum = 00:03:27<br>Maximum = 00:44:36 |
| **Dragonflye hybrid** | Hybrid | Median = 00:04:34<br>Minimum = 00:01:32<br>Maximum = 00:07:27 | Median = 00:03:46<br>Minimum = 00:01:22<br>Maximum = 00:06:01 |
| **Unicycler** | Hybrid | Median = 00:50:25<br>Minimum = 00:12:04<br>Maximum = 01:13:32 | Median = 00:34:10<br>Minimum = 00:08:36<br>Maximum = 00:48:23 |
| **Hybracter long** | Long | Median = 00:11:46<br>Minimum = 00:03:26<br>Maximum = 00:36:09 | Median = 00:10:20<br>Minimum = 00:03:17<br>Maximum = 00:29:50 |
| **Dragonflye long** | Long | Median = 00:04:10<br>Minimum = 00:01:22<br>Maximum = 00:06:01 | Median = 00:04:34<br>Minimum = 00:01:32<br>Maximum = 00:07:27 |

The difference in runtime performance between Hybracter and Dragonflye is predominantly the result of the included targeted plasmid assembly and the reorientation and assessment steps in Hybracter that are not included in Dragonflye. Additionally, the results suggest limited benefits to running Hybracter with more than eight threads. As explained in the following section, if a user has multiple isolates to assemble, a superior approach is to modify the configuration file specifying more efficient resource requirements for each job in Hybracter.

### Parallelisation Allows for Improved Efficiency

Hybracter allows users to specify and customise a configuration file to maximise resource usage and runtime efficiency. Users can modify the desired threads, memory and time requirements for each type of job that is run within Hybracter to suit their computational resources. So that resources are not idle for most users on single sample assemblies, large jobs such as the Flye and Plassembler assembly steps default to 16 threads and 32 GB of memory.

To emphasise the efficiency benefits of parallelisation, the 12 *Lerminiaux* et al. isolates were also assembled using ‘hybracter hybrid’ with a customised configuration file designed to improve efficiency on the machine used for benchmarking. Specifically, the configuration was changed to specify 8 threads and 16 GB of memory allocated to big jobs (assembly, polishing and assessment) and 4 threads and 8 GB of memory allocated to medium jobs (reorientation). More details on changing Hybracter’s configuration file to suit specific systems can be found in the documentation (https://hybracter.readthedocs.io/en/latest/configuration/). We limited the overall ‘hybracter hybrid’ run with 32 GB of memory and 16 threads to provide a fair comparison. The overall ‘hybracter hybrid’ run was then compared to the sum of the 12 ‘hybracter hybrid-single’ runs. Overall, the 12 isolates took 01h48m57s in the combined run, as opposed to 04h38m45s from the sum of the 12 ‘hybracter hybrid-single’ and 07h04m04s from the sum of the 12 Unicycler runs. This inbuilt parallelisation of Hybracter provides significant efficiency benefits if multiple samples are assembled simultaneously. The performance benefit of Hybracter afforded by Snakemake integration in parallel computing systems may be variable over different architectures, but this provides an example case of potential efficiency and convenience benefits.

### Long-Read Depth Does Not Affect Hybrid Assembly Accuracy If a Complete Chromosome is Assembled

Finally, we tested the effect of long-read depth on the accuracy of assemblies with all five tools at an estimated long-read depth from 10x to 100x at every interval of 5x for an example isolate (*Lerminiaux* Isolate B, *Enterobacter cloacae*) with super-accuracy model basecalled simplex reads (Figure 4 and Supplementary Table 12). At 10x and 15x sequencing depth, only Unicycler was able to assemble a complete chromosome. From 20x and above, all five tools were able to assemble complete chromosomes. For the hybrid tools, once a complete chromosome was assembled, increasing long-read depth had a negligible impact on accuracy results (Figure 4). Notably, Hybracter hybrid was able to produce perfect assemblies from as low as 20x long-read depth. For long-read only tools, increasing long-read depth did affect accuracy. Increasing depth improved SNV accuracy for both Hybracter long and Dragonflye long (Figure 4C). For small InDels, Hybracter long improved with extra depth, while Dragonflye long actually performed worse (Figure 4A). Depth had minimal impact on large InDels (Figure 4B).

**Figure 4:**
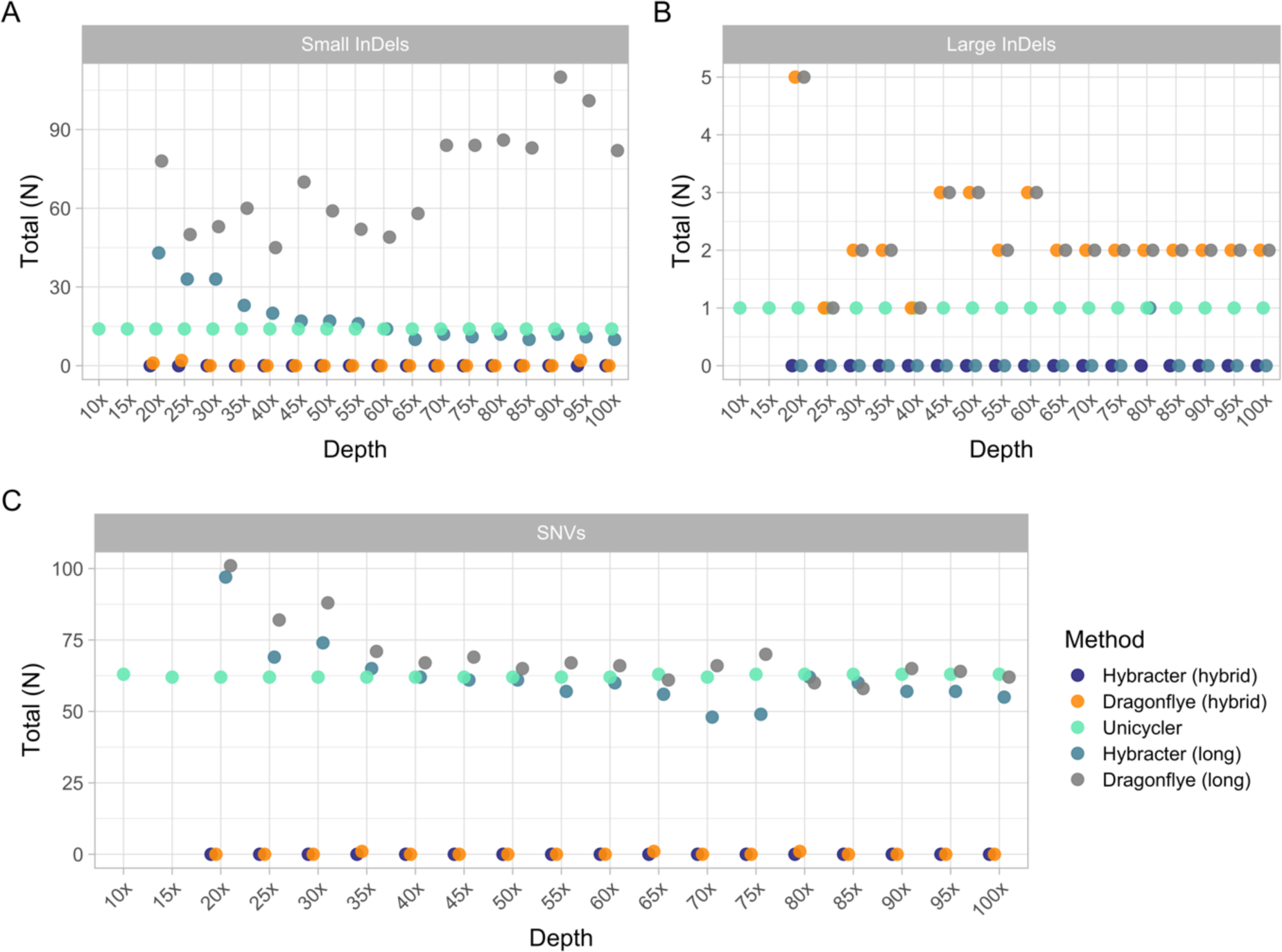
Comparison of the counts of small (<60bp) (A) and large (>60bp) (B) insertions and deletions (InDels) and SNVs (C) for Hybracter hybrid, Dragonflye hybrid, Unicycler, Hybracter long and Dragonflye long chromosome assemblies of *Lerminiaux* Isolate B (*Enterobacter cloacae*) at 5x intervals of sequencing depth from 10x to 100x.

## Discussion

As long-read sequencing has improved in accuracy with reduced costs, it is now routine to use a combination of long- and short-reads to generate complete bacterial genomes^3,5^. Recent advances in assembly algorithms and accuracy improvements mean that a long-read first hybrid assembly should be favoured with short-reads being used after assembly for polishing ^12^, as opposed to the short-read first assembly approach (where long-reads are only used for scaffolding a short-read assembly) utilised by the current gold standard automated assembler Unicycler. The Unicycler approach is more prone to larger scale InDel errors as well as smaller scale errors such as those caused by homopolymers or methylation motifs ^6,11,52,53^. Additionally, it should be noted that it is already possible (while perhaps not routine) to generate perfect hybrid bacterial genome assemblies using manual consensus approaches requiring human intervention, such as Trycycler^7,54^. While manual approaches such as Trycycler generally yield superior results to automated approaches, manually assembling many complete genomes is challenging as considerable time, resources and bioinformatics expertise are required.

The results of this study emphasise that the long-read first hybrid approach consistently yields superior assemblies than the short-read first hybrid approach and should therefore be preferred going forward. The only exception where a short-read first approach is to be preferred is where a limited depth of long-read sequencing data is available (<20x depth). In this instance, long-read first hybrid approaches may struggle to assemble a complete chromosome, while short-read first approaches like Unicycler may be able to (Figure 4).

Interestingly, in the course of conducting benchmarking for this study, we found a large number of discrepancies between older short-read first assembled ‘reference genomes’ for *S. aureus* JKD6159 ^55^ and the five ATCC genomes benchmarked compared to updated Trycycler long-read first references (See Supplementary Table 13). The number of discrepancies ranged from 44 to 8,255 across the six genomes. Therefore, we recommend that older short-read first reference genomes be updated if possible using a long-read assembly approach (such as with Trycycler).

This study also shows that automated perfect hybrid genome assemblies are already possible with Hybracter. This study and others^9,54^ also confirm that a long-read first hybrid approach remains preferable to long-read only assembly with Nanopore reads, as short-reads continue to provide accuracy improvements in polishing steps. However, it is foreseeable that short-reads will soon provide little or no accuracy improvements and will not be needed to polish long-read only assemblies to perfection. Already, perfect long-read only assemblies are possible, at least with manual intervention using Trycycler ^7^. Accordingly, automated perfect bacterial genome assemblies may soon become possible from long-reads only. Hybracter also allows users to turn long-read polishing off altogether. It is already established that long-read polishing can introduce errors and make long-read only assemblies worse with highly accurate Nanopore and PacBio reads^11,31^. Therefore, this feature may become increasingly useful as long-read sequencing continues to improve in accuracy and we recommend its use for highly accurate Q20+ long-reads.

Hybracter was created to bridge the gap from the present to the future of automated perfect hybrid and long-read-only bacterial genome assemblies. The results of this study show that Hybracter in hybrid mode is both faster and more accurate than the current gold standard tool for hybrid assembly Unicycler and is more accurate than Dragonflye in both modes. It should be noted that if users want fast chromosome-only assemblies where accuracy is not essential (for applications such as species identification or sequence typing), Dragonflye remains a good option due to its speed.

Hybracter especially excels in recovering complete plasmid genomes compared to other tools. By incorporating Plassembler, Hybracter recovers more complete plasmid genomes than Unicycler in hybrid mode. Further, Hybracter long is comparable to Unicycler and Hybracter hybrid when using long-reads only for plasmid recovery.

The high error rates of long-read sequencing technologies have prevented the application of assembly approaches designed for highly accurate short-reads, such as constructing de Bruijn graphs (DBGs) based on strings of a particular length *k* (*k*-mers) ^56–58^. This resulted in bioinformaticians initially utilising less efficient algorithms designed with long-reads in mind, such as utilising overlap graphs in place of DBGs^27,37,39,59,60^. While DBGs have been used for long-read assembly in some applications ^61–63^, adoption, especially in microbial genomics, has been limited.

Although long-read first assembly methods enable complete chromosome and large plasmid reconstruction, it is well established that long-read only assemblers struggle to assemble small (<20kbp) plasmids accurately, often leading to missing or multiplicated assemblies^6,51^. These errors may be exacerbated if ligation chemistry-based sequencing kits are used^51^. Therefore, hybrid DBG based short-read first assemblies are traditionally recommended for plasmid recovery^12^.

Implemented in our post-publication changes to Plassembler described in this study, Hybracter solves the problem of small plasmid recovery using long-reads. It achieves this by implementing a DBG-based assembly approach with Unicycler. The same read set is used twice, first as unpaired pseudo ‘short’ reads and then as long-reads; the long-read set scaffolds a DBG-based assembly based on the same read set. This study demonstrates that current long-read technologies, such as R10 Nanopore reads, are now accurate enough that some short-read algorithms are applicable. Our results also suggest that similar DBG-based algorithmic approaches could be used to enhance the recovery of small replicons in long-read datasets beyond the use case presented here of plasmids in bacterial isolate assemblies. This could potentially enhance the recovery of replicons such as bacteriophages^64^ or other small contigs from metagenomes using only long-reads^10,50^.

Finally, consistent and resource efficient assemblies that are as accurate as possible in recovering both plasmids and chromosomes are crucial, particularly for larger studies investigating plasmid epidemiology and evolution. AMR genes carried on plasmids can have complicated patterns of transmission involving horizontal transfer between different bacterial species and lineages, transfer between different plasmid backbones, and integration into and excision from the bacterial chromosome^65–67^. Accurate plasmid assemblies are crucial in genomic epidemiology studies investigating transmission of antimicrobial resistant bacteria within outbreak settings, as well as in a broader One Health context, where hundreds or even thousands of assemblies may be analysed^68–71^. Hybracter will facilitate the expansion of such studies, allowing for faster and more accurate automated complete genome assemblies than existing tools. Additionally, by utilising Snakemake^20^ with a Snaketool^21^ command line interface, Hybracter is easily and efficiently parallelised to optimise available resources over various large-scale computing architectures. Individual jobs (such as each assembly, reorientation, polishing or assessment step) within Hybracter are automatically sent to different resources on a high performance computing (HPC) cluster using the HPC’s job scheduling system like Slurm^72^. Hybracter can natively use any Snakemake-supported cloud-based deployments such as Kubernetes, Google Cloud Life Sciences, Tibanna, and Azure Batch.

## Conclusion

Hybracter is substantially faster than the current gold standard automated tool Unicycler, assembles chromosomes more accurately than existing methods, and is superior at recovering complete plasmid genomes. By applying DBG-based algorithms designed for short-reads on current generation long-reads, Hybracter long also solves the problem of long-read-only assemblers entirely missing or duplicating small circular elements such as plasmids. Hybracter is resource efficient and natively supports deployment on high-performance computer clusters and cloud environments for massively parallel analyses. We believe Hybracter will prove to be an extremely useful tool for the automated recovery of complete bacterial genomes from hybrid and long-read-only sequencing data suitable for massive datasets.

## Acknowledgements

This work was supported with supercomputing resources provided by the Phoenix HPC service at the University of Adelaide. We would particularly like to thank Fabien Voisin for his integral role in maintaining and running Phoenix. We would also like to thank Brad Hart for useful comments in testing Hybracter and Simone Pignotti, Yu Wan and Oliver Schwengers for providing helpful comments and GitHub pull requests.

---

**Supplementary Figure 1:**
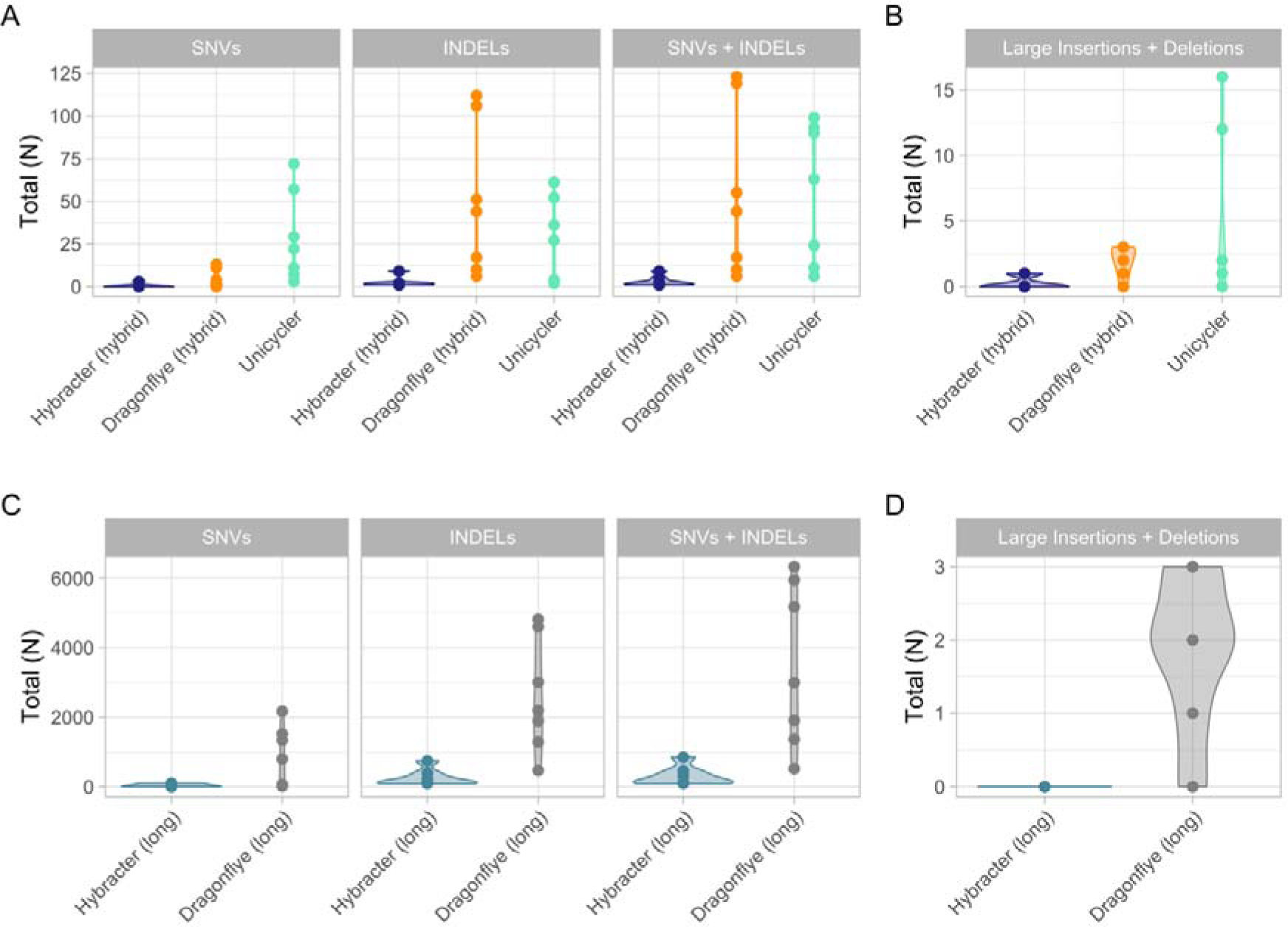
For lower quality samples only (all five ATCC fast model base-called samples, *S. aureus* JKD6159 with R9 chemistry and *M.* tuberculosis H37R2), this figure presents a comparison of the counts of small nucleotide variants (SNVs) and small (<60bp) insertions and deletions (InDels) (A) and the total number of large (>60bp) InDels (B) for the hybrid tools benchmarked (Hybracter hybrid in dark blue, Dragonflye hybrid in orange and Unicycler in green). The counts of SNVs and small InDels (C) and the total number of large InDels (D) for the long tools benchmarked (Hybracter long in light blue, Dragonflye long in grey) are also shown. All data presented is from the benchmarking output run with 8 threads.

**Supplementary Figure 2:**
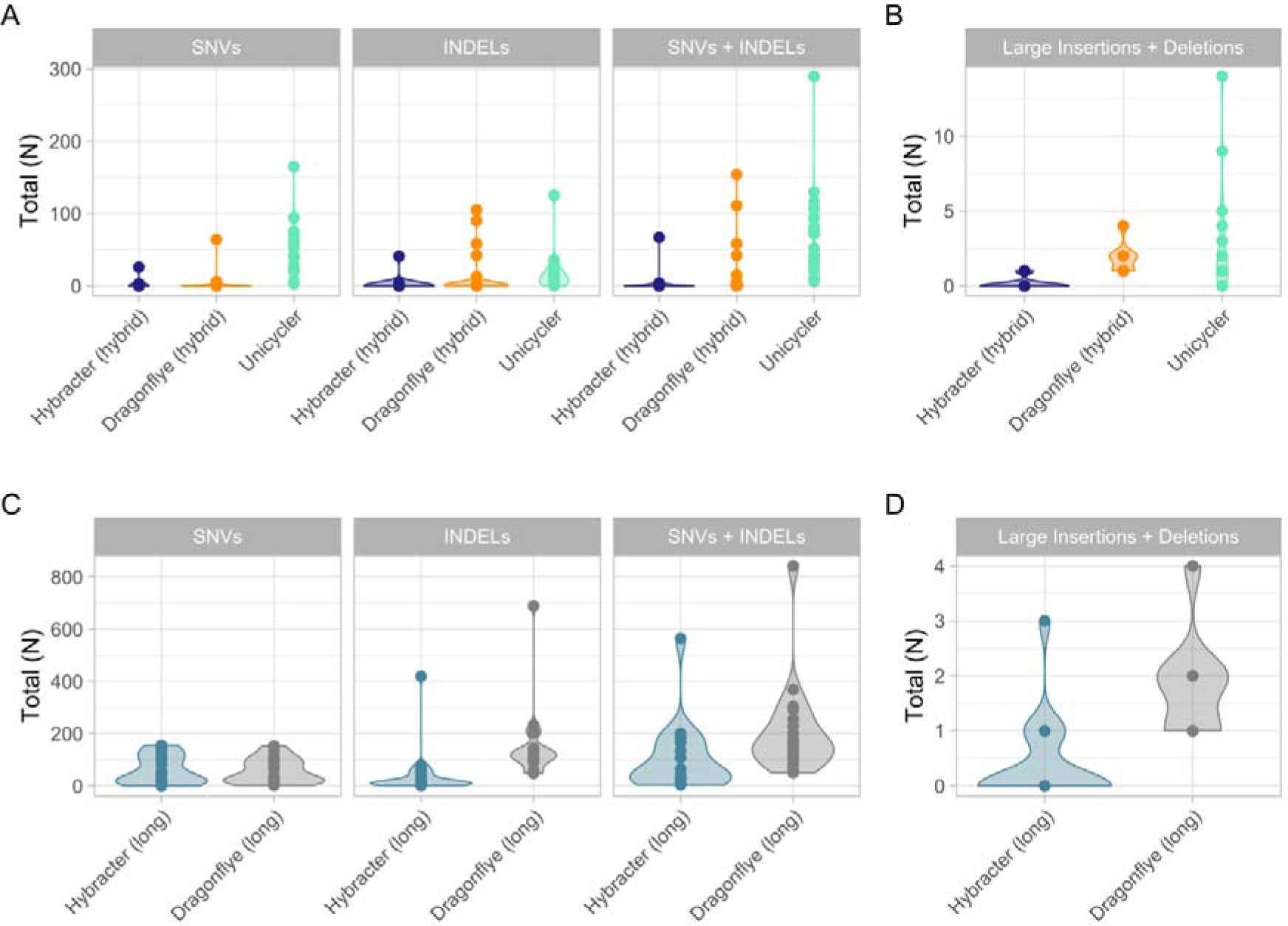
For simplex super-accuracy model basecalled samples (five ATCC, *S. aureus* JKD6159 with R10 chemistry and all twelve *Lerminiaux* et al. samples), this figure presents a comparison of the counts of small nucleotide variants (SNVs) and small (<60bp) insertions and deletions (InDels) (A) and the total number of large (>60bp) InDels (B) for the hybrid tools benchmarked (Hybracter hybrid in dark blue, Dragonflye hybrid in orange and Unicycler in green). The counts of SNVs and small InDels (C) and the total number of large InDels (D) for the long tools benchmarked (Hybracter long in light blue, Dragonflye long in grey) are also shown. All data presented is from the benchmarking output run with 8 threads.

**Supplementary Figure 3:**
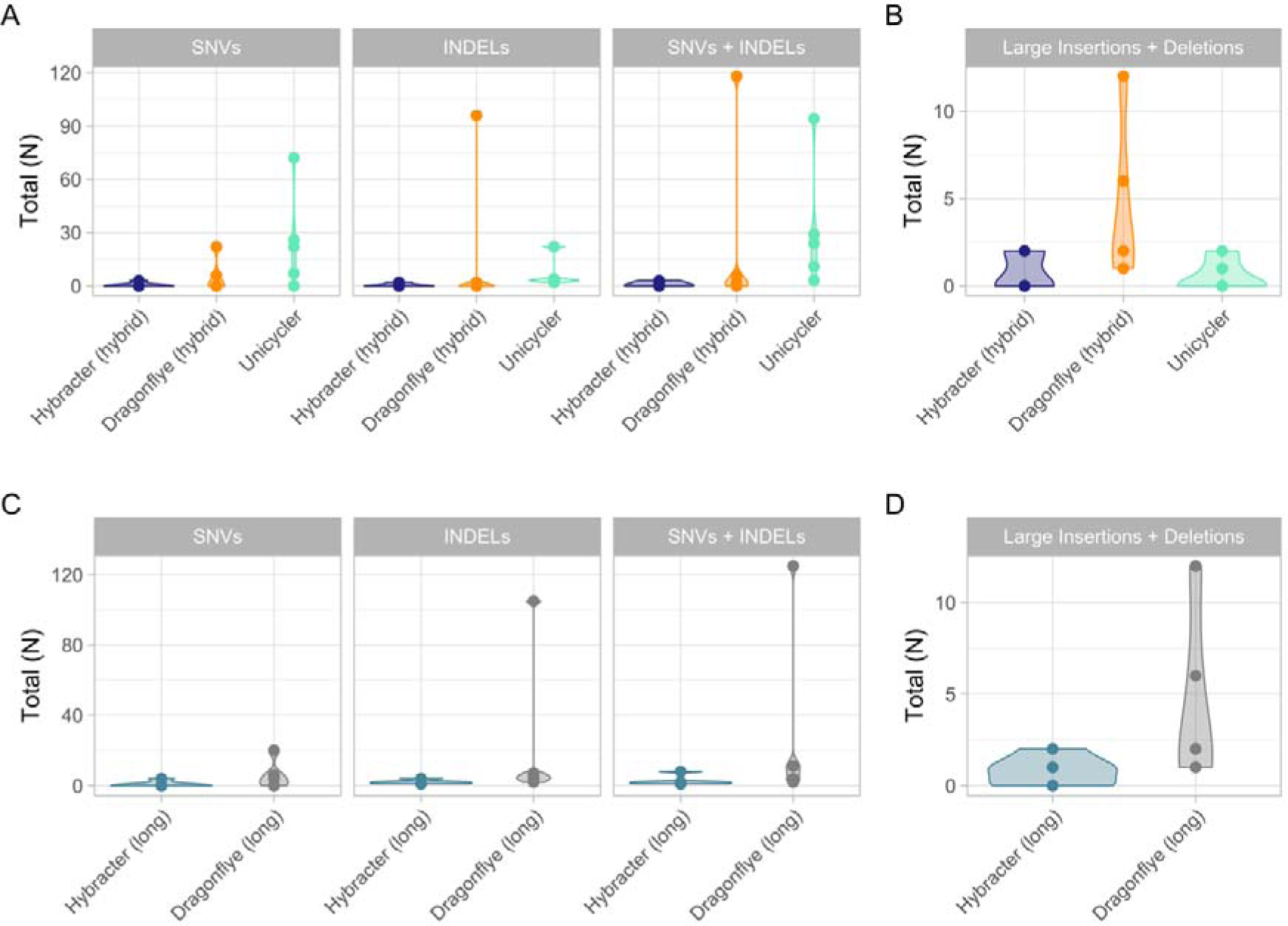
For duplex super-accuracy model basecalled samples (five ATCC samples), this figure presents a comparison of the counts of small nucleotide variants (SNVs) and small (<60bp) insertions and deletions (InDels) (A) and the total number of large (>60bp) InDels (B) for the hybrid tools benchmarked (Hybracter hybrid in dark blue, Dragonflye hybrid in orange and Unicycler in green). The counts of SNVs and small InDels (C) and the total number of large InDels (D) for the long tools benchmarked (Hybracter long in light blue, Dragonflye long in grey) are also shown. All data presented is from the benchmarking output run with 8 threads.

